# Shared and distinct patterns of atypical cortical morphometry in children with autism and anxiety

**DOI:** 10.1101/2020.03.26.010272

**Authors:** Shelly Yin, Seok-Jun Hong, Adriana Di Martino, Michael P. Milham, Bo-Yong Park, Oualid Benkarim, Richard A.I. Bethlehem, Boris C. Bernhardt, Casey Paquola

**Author notes:** shared co-authorship (Boris C. Bernhardt, Casey Paquola).

## Abstract

Autism spectrum disorder (ASD) and anxiety disorders (ANX) are common neurodevelopmental conditions with several overlapping symptoms. Notably, many children and adolescents with ASD also have an ANX diagnosis, suggesting shared pathological mechanisms. Here, we leveraged structural imaging and phenotypic data from 82 closely matched children (28 ASD, 28 ANX, 26 typically developing controls). Our neuroimaging paradigm assessed cortical thickness and studied inter-regional structural covariance networks. Both ASD and ANX presented with atypical structural network organization relative to controls. Specifically, ASD presented with increased thickness in temporal and parietal midline cortices, while ANX was associated with increased cortical thickness in the left inferior frontal and precentral gyri. Despite the overall difference in the spatial distributions these clusters, unconstrained spin permutation testing showed that statistical maps from the ANX-vs-controls and ASD-vs-controls analyses were significantly correlated. The two diagnostic groups also presented with common decreases in structural covariance patterns, collectively pointing to decreased structural coupling between lateral frontal, lateral temporal, and temporo-parietal regions. Dimensional analysis of trait anxiety and social responsiveness partially recapitulated diagnosis-based findings. Collectively, our findings provide evidence for both shared as well as distinct effects of ASD and ANX on on regional and inter-regional structural network organization.

## 1. INTRODUCTION

Autism spectrum disorder (ASD) and anxiety disorders (ANX) are two of the most prevalent neuropsychiatric conditions affecting young people (Blumberg et al., 2013; Krain et al., 2007; Twenge et al., 2009; Xu et al., 2009) that typically persist into adulthood (Kessler et al., 2005; Mandell, Novak, and Zubritsky, 2005). Traditionally, both are diagnosed based on clinical history and symptomatology (Mullin et al., 2013), and their study has provided valuable human evidence on different aspects of social and affective processes. ASD has frequently been associated with atypical social cognition (Baron-Cohen, 1997; Frith and Happe, 1994), while ANX relate to atypical emotional reactivity and regulation (Campbell-Sills and Barlow, 2007; Cisler et al., 2010). Despite this conceptual distinction, high comorbidities of ASD and ANX render the situation more difficult, with 40% of children and adolescents with ASD having a concurrent ANX diagnosis (van Steensel et al., 2011), suggesting the existence of common neurodevelopmental perturbations (White et al., 2009; van Steensel et al., 2011). High comorbidity rates may also be due to challenges in differential diagnosis using current measures, which adds impetus to studying ASD and anxiety in a dimensional manner. A handful of neuroimaging studies have demonstrated qualitatively distinct effects of anxiety and ASD on amygdala volume and task-related activations (Herrington, Maddox, Kerns, *et al.*, 2017; Herrington, Maddox, McVey, *et al.*, 2017; Ibrahim *et al.*, 2019), however, we currently have a poor understanding of the shared and unique effects on the structure of the cortex.

Magnetic resonance imaging (MRI) is widely accessible and enables quantification of brain morphology, making it a good candidate to identify intermediary *in vivo* phenotypes of complex mental health conditions such as ASD and ANX. One of the most widely used markers is cortical thickness, measured as the distance between inner and outer cortical interfaces on structural MRI. Cortical thickness provides biologically meaningful insight regarding the cellular and synaptic organization of the cortex (Desrivieres et al., 2014; Huttenlocher, 1979; Schuz and Palm, 1989). Previous studies have reported varied patterns and directions of cortical anomalies in ASD relative to typically developing controls (TDC) (Khundrakpam et al., 2017; Redcay, 2007; Pereira et al., 2018; Raznahan et al., 2010; Scheel et al., 2011; Wallace et al., 2010); however, recent large-scale studies based on open datasets have generally converged on increased thickness in frontal and temporal cortical areas in individuals with ASD (Bedford et al., 2019; Hong et al., 2017; Valk et al., 2015). Studies assessing cortical morphology in ANX have also pointed to increased cortical thickness in medial and lateral frontal regions relative to TDC (Gold et al., 2017; Strawn et al., 2014), with, however, a seemingly different spatial topography compared to ASD. It thus remains to be established whether syndromic differences are also reflected in divergent signatures of regional morphology.

In addition to mapping effects of specific brain regions, cortical thickness measures can also be harnessed via covariance paradigms to inspect inter-regional coordination of different brain areas, allowing evaluation of structural network organization (Alexander-Bloch et al., 2013; Lerch et al., 2006; Mechelli et al., 2005). Applied extensively to cohorts of children and adolescents, structural covariance analysis may approximate common maturation patterns (Zielinski et al., 2010) and partially recapitulates intrinsic functional networks (Seeley et al., 2009). Furthermore, work in typically developing individuals and those with neuropsychiatric disorders has shown that inter-individual differences in socio-cognitive and affective capacities can be captured by variations in structural covariance, even when these phenotypic measures do not relate to regional variations in brain structure per se (Bernhardt et al., 2014a,b; Valk et al., 2017). Therefore, cortical thickness and covariance analyses can offer complementary information on the coupling of macroscale structural network organization with ASD- and ANX-related behaviors.

The current work investigated shared and distinct structural substrates of ASD and ANX in the cortex, providing an *in vivo* neuroanatomical complement to previous clinical and pharmacological studies (van Steensel et al., 2011; Vasa and Mazurek, 2015). The importance of community-representative cohorts for translation and inclusivity in psychiatric research has been clearly asserted for clinical trials (Geddes, 2005; Surman *et al.*, 2010), but is less acknowledged in neuroimaging. Based on these recommendations, ANX and ADHD co-morbidities were nested within the clinical samples, and we used categorical and dimensional analyses to strike a trade-off between interpretability and external validity (Kraemer et al., 2004; Kapur et al., 2012). Categorical analyses inform upon the common abnormalities within a primary diagnosis, whereas dimensional analyses illustrate the relevance of cortical malformations to a specific clinical symptom. In light of the comorbidity of ANX in ASD, we predicted there to be shared morphological alterations in both primary diagnostic groups compared to controls. We nevertheless also hypothesized that cortical thickness differences would be concentrated within functional networks relevant to disorder-specific deficits, and these would overlap with dimensional associations of core symptoms. Specifically, we expected more marked structural alterations in ANX within networks previously implicated in salience and emotion processing (Menon, 2015; Seeley et al., 2007), and changes related to ASD within networks that may more generally contribute to socio-cognitive processing, such as the default mode network (Schilbach et al., 2008). Our work leveraged a first wave of data provided by the Healthy Brain Network (HBN), an ongoing and large-scale transdiagnostic sample aggregating imaging and phenotypic data in typically developing children and adolescents as well as individuals with a neuropsychiatric diagnosis, that allowed directed comparison across the ASD, ANX and TDC cohorts (Alexander et al., 2017).

## 2. MATERIALS AND METHODS

### 2.1 Participants

We studied the open-access Child Mind Institute HBN dataset (Alexander et al., 2017), releases 1-4 collected between June 2017 and July 2018. The dataset aims to represent a broad range of heterogeneity in developmental psychopathology. Participants were recruited using a community-referral model (for inclusion criteria, see http://fcon_1000.projects.nitrc.org/indi/cmi_healthy_brain_network/inclusion.html). HBN was approved by the Chesapeake Institutional Review Board. Written informed consent was obtained from all participants and from legal guardians of participants younger than 18 years.

The HBN protocol consists of four 3-hour sessions collecting general information, behavioral measures, diagnostic assessments, and neuroimaging data (for a complete list of measures, please see http://fcon_1000.projects.nitrc.org/indi/cmi_healthy_brain_network/assessments.html). Psychiatric diagnoses were assessed and reported by clinicians according to DSM-5 criteria. Among the 576 individuals from releases 1-4 with T1-weighted MRI available, we restricted inclusion to participants with no diagnosis and those diagnosed with ASD and/or ANX. Participants were categorized by primary diagnosis. Of note, all participants with ASD and ANX presented with a primary diagnosis of ASD. Exclusion criteria were any other psychiatric or intellectual comorbidites, except for ADHD in the ASD and ANX groups. We chose to include participants with a secondary diagnosis of ADHD in these groups due to the high prevalence and to provide a more representative community sample of the diagnoses. Although 3 collection sites provided data to the included releases of the HBN dataset, we further restricted inclusion to participants from the Rutgers University Brain Imaging Centre (RUBIC) as the other sites (Staten Island, Citigroup Biomedical Imaging Centre) were inadequately represented in each group in these initial releases. This initial cohort thus consisted of 172 participants: 41 ASD (10.78±3.91 years, 12 females), 81 ANX (11.52±3.72 years, 36 females), 50 TDC (10.12±3.61 years, 26 females).

### 2.2 MRI acquisition

Imaging at the RUBIC was conducted using a Siemens 3T Tim Trio scanner with a 32-channel head coil. 3D T1-weighted sagittal magnetization-prepared rapid acquisition gradient echo (MPRAGE) structural images were obtained with the following parameters: repetition time (TR) = 2500 ms, echo time (TE) = 3.15 ms, flip-angle = 8, Field-of-View (FoV) = 256 mm^2^, resulting in 224 slices with 0.8 × 0.8 × 0.8 mm^3^ voxels (Alexander et al., 2017).

### 2.3 Imaging processing and quality control

FreeSurfer (v6.0; http://surfer.nmr.mgh.harvard.edu) was used to generate cortical surface models and to measure cortical thickness (Fischl and Dale, 2000). In brief, FreeSurfer automatically reconstructs geometric models of inner and outer cortical interfaces using a series of volume- and surface-based processing steps. Extracted surfaces in each individual were registered to fsaverage5, an average spherical representation with 20,484 surface points, by aligning cortical folding patterns. Surface extractions were visually inspected and segmentation inaccuracies were manually corrected by one rater (SY) blinded to participant diagnoses. Thickness data were smoothed using a 20 mm Gaussian kernel. This process reduces noise and misalignment between vertices by replacing values in images as a weighted average of itself and its neighboring vertices (Lerch and Evans, 2005).

### 2.4 Participant matching

We excluded 56 participants because of poor image quality due to excessive head motion or low tissue contrast. The matchit package in R (v3.2.5; https://cran.r-project.org/web/packages/MatchIt/MatchIt.pdf) was used to match the remaining participants across groups on age and sex as well as ADHD comorbidity for ASD and ANX groups, in order to reduce model dependence and potential for bias (Ho *et al.*, 2007). The argument specifications were “nearest” for method with a ratio of 1, indicating that participants should be matched as closely as possible and only once. The final cohort used in structural analysis consisted of 82 participants: 28 ASD (11.46±3.96 years, 8 females), 28 ANX (12.07±3.48 years, 8 females), and 26 TDC (12.08±3.43 years, 14 females). **Tables 1–2** summarize changes in the groups before and after matching, as well as the final cohort.

**Table 1.** Balance of age, sex, and comorbidity of ADHD before/after matching.

|  | Unmatched | Matched | Balance Improvement (%) |
| --- | --- | --- | --- |
| *Age* |  |  |  |
| ASD | 11.46 | 11.46 | 38 |
| ANX | 12.45 | 12.07 |  |
| Mean Difference | 0.98 | 0.61 |  |
| *Sex (females)* |  |  |  |
| ASD | 8 | 8 | 100 |
| ANX | 30 | 8 |  |
| Mean Difference | 0.23 | 0.00 |  |
| *ADHD* |  |  |  |
| ASD | 18 | 18 | 100 |
| ANX | 26 | 18 |  |
| Mean Difference | 0.19 | 0.00 |  |

**Table 2.** Demographic and phenotypic information of participants with a primary diagnosis of autism spectrum disorder (ASD) or anxiety disorder (ANX), or typically developing controls (TDC).

|  | ASD<br>(n=28) | ANX<br>(n=28) | TDC<br>(n=26) | Group difference | Significant post-hoc tests |
| --- | --- | --- | --- | --- | --- |
| Age | 11.46 ± 3.96 | 12.07 ± 3.48 | 12.08 ± 3.43 | F(80)=3.24, p=0.72 |  |
| Sex (females) | 8 | 8 | 14 | F(80)=2.16, p=0.12 |  |
| ADHD diagnosis | 18 | 18 | 0 | $\chi^2=31.4$, p<0.01 | TDC<ASD<br>TDC<ANX |
| ANX diagnosis | 8 | 28 | 0 | $\chi^2=58.8$, p<0.01 | TDC<ANX<br>ASD<ANX |
| WISC<sup>1</sup> | 94.9 ± 16.5 | 98.9 ± 16.5 | 108.0 ± 11.7 | F(39)=2.44, p=0.10 |  |
| SCARED score<sup>2</sup> | 22.98 ± 13.33 | 26.30 ± 10.96 | 13.12 ± 7.34 | F(66)=8.84, p<0.01 | TDC<ASD<br>TDC<ANX |
| SRS-2 score<sup>3</sup> | 91.42 ± 26.87 | 62.00 ± 26.15 | 29.17 ± 24.09 | F(73)=11, p<0.01 | TDC<ANX<ASD |

<sup>1</sup> Only available for n=41<sup>2</sup> Only available for n=68<sup>3</sup> Only available for n=75

### 2.5 Phenotypic assessments

In addition to analyzing the effects of dichotomous ASD and ANX diagnoses, we implemented a dimensional approach based on continuous phenotypes reflecting overall symptoms. We focused on the Social Responsiveness Scale (SRS-2) and the Screen for Child Anxiety Related Disorders (SCARED) to index autism and anxiety risk, respectively. Both scales have moderate-to-high internal consistency, interrater reliability, and test-retest reliability (Bölte et al., 2008; Bruni, 2014; Su et al., 2008). SRS-2 measures characteristic deficits of social interaction and communication in ASD and consists of 65 items rated on a 3-point scale by parents (Constantino et al., 2003) of participants ages 5-17 years. SCARED is a questionnaire consisting of 41 items rated on a 3-point scale that screens for childhood anxiety (Birmaher et al., 1999) and was administered to parents of and participants aged 8-17 years. Parent- and self-report components of SCARED were moderately correlated (r=0.41, *p*<0.001); in line with prior work (Gold et al., 2017; Ivarsson et al., 2018), a combined score was used. Completed SRS-2 data were available for 75/82 participants (24 ASD, 27 ANX, 24 TDC) and SCARED data for 68/82 (22 ASD, 25 ANX, 21 TDC) (Table 2).

### 2.6 Statistical Analysis

Surface-based analysis was carried out using SurfStat (https://mica-mni.github.io/surfstat/; Worsley et al. 1999) for Matlab (R2017b, The Mathworks, Natick, MA). All models accounted for sex and age.

#### a) Cortical thickness comparisons

Linear models compared cortical thickness at each vertex *i* between ANX and TDC as well as ASD and TDC. We also directly compared ASD and ANX. The corresponding model at each vertex was:

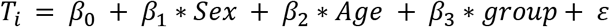

To assess shared substrates, we furthermore computed Pearson correlations between the t-statistic maps of the above contrast. Spatial dependencies are produced in cortical measurements by smoothing and motion artefacts, as well as the spatial constraints of brain organisation. Parametric tests on the correspondence of spatial maps falsely assume spatial independence, however, leading to high false positive rates. Instead we determined the significance of spatial map correspondence using the spatial spin permutation test method with 1000 permutations (Alexander-Bloch et al., 2018; Vos de Wael et al., 2019). In brief, this method generates a null distribution by comparing a spatial map to a permutated map created by applying random rotational permutations to a spherical representation of a cortical surface (Alexander-Bloch et al., 2018).

#### b) Structural covariance analysis

We carried out a structural covariance analysis centered on clusters of significant cortical thickness differences identified in *a)*. Analyses were first conducted within TDC to map typical covariance networks. The model fitted for a given cluster was:

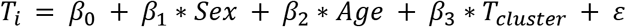

To assess between-group differences in covariance, we fitted linear interaction models as in previous work (Bernhardt et al., 2008; Bernhardt et al., 2014b).

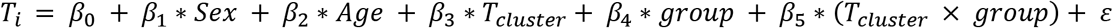

#### c) Dimensional analysis

We compared cortical thickness at each vertex across all participants to assess the effect of SRS-2 or SCARED score with the model:

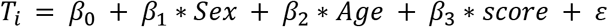

We also repeated structural covariance analyses with SRS-2 or SCARED score, after regressing out the effects of sex and age (with or without additional control for diagnosis group). To reveal structural covariance effects, we furthermore stratified participants as either high or low scoring on the phenotypic assessments using a median split.

#### d) Correction for multiple comparisons

Findings were corrected for multiple comparisons using random field theory for non-isotropic images (Worsley et al. 1999), unless otherwise stated. This controlled the chance of reporting a family-wise error (FWE) to p<0.05. As in previous work, a cluster defining threshold of p<0.025 was used (Valk et al., 2017).

## 3. RESULTS

### 3.1 Shared and distinct cortical thickness signatures of ASD and ANX

Surface-based cortical thickness mapping showed significant differences of moderate effect sizes between each pair of groups. We observed increased thickness in ASD compared to TDC in the right middle temporal regions (*MTG*; t=2.54, p_FWE_=0.002, Cohen’s d=0.74) and right retrosplenial cortex (*RSC*; **Figure 1A**; t=2.62, p_FWE_=0.003, d=0.50). Relative to TDC, ANX was related to increased thickness in left inferior frontal and precentral gyri (*IFG*; **Figure 1A**; t=2.54, p_FWE_=0.003, d=0.59). We detected moderate spatial correspondence of the ASD-vs-TDC and ANX-vs-TDC t-statistic maps (r=0.49), and spin permutation tests indicated that this similarity was unlikely attributable to shared spatial autocorrelation alone (**Figure 1C**; p_spin_<0.001). Directly comparing the clinical cohorts, ASD showed increased thickness in the cuneus (*CUN*; **Figure 1B**;t=2.49, p_FWE_=0.007, d=0.46).

**Figure 1.**
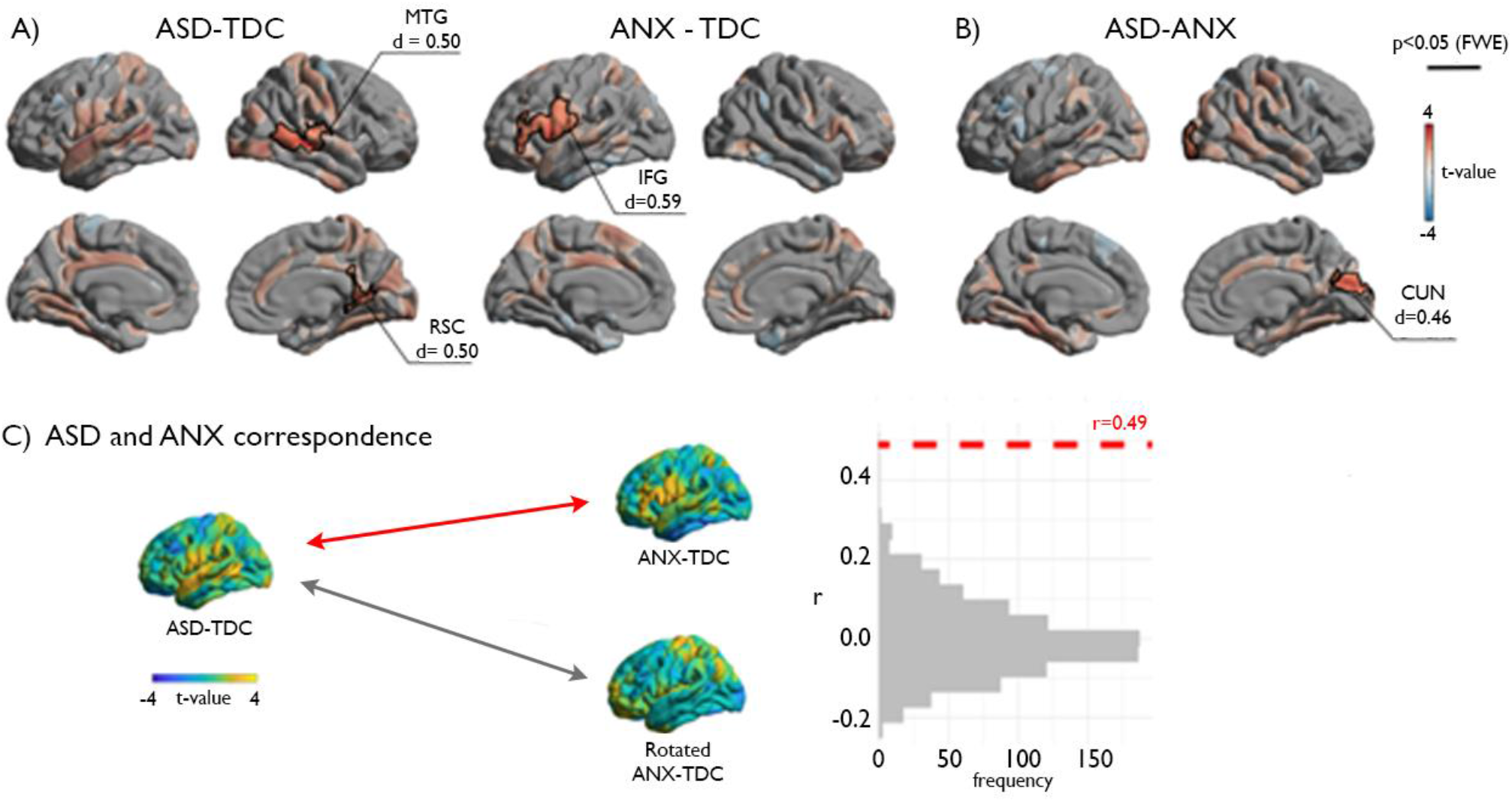
Cortical thickness analysis. **(A, B)** Cortical thickness comparisons between the different cohorts (ASD – autism spectrum disorder, ANX – anxiety, TDC – typically developing controls). Red/blue indicates increases/decreases in cortical thickness based on the relative contrast. Trend level differences are semi-transparent and findings corrected for multiple corrections are outlined in black. The average Cohen’s d effect size of each significant cluster is indicated. (**C**) Spatial map correspondence between ASD-vs-TDC and ANX-vs-TDC, indicated in red, compared to a null model distribution generated by the spin permutation method.

#### 3.2 Covariance network modulations

Focusing on regions of significant between-group differences in cortical thickness (*Figure 1*), we evaluated large-scale morphological networks using seed-based covariance analyses. Covariance patterns for all clusters were first mapped within TDC to visualize typical networks, followed by interaction models to assess between-group differences. For all clusters, covariance networks were not solely limited to the ipsilateral perimeter of the cluster but encompassed bilaterally distributed networks. Specifically, the IFG showed widespread covariance to bilateral prefrontal, fronto-central, midline parietal-occipital, and right lateral occipital regions (**Figure 2B**). Compared to TDC, ANX diagnosis was associated with weaker covariance of the IFG to ipsilateral inferior temporal (p_FWE_=0.007) and lateral occipital (p_FWE_=0.02) as well as bilateral temporo-parietal regions (p_FWE_<0.005) (**Figure 2B, Table 3**).

**Figure 2.**
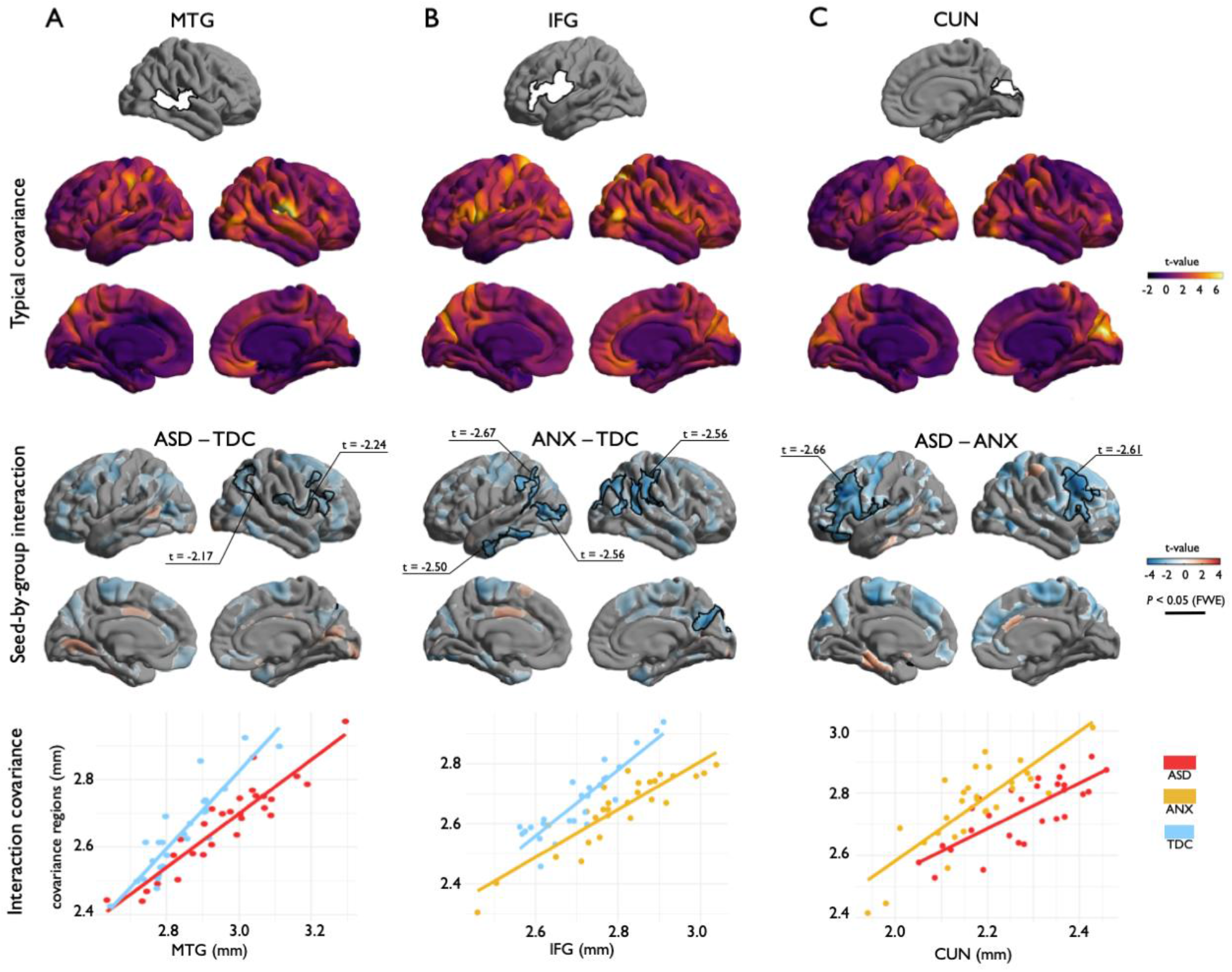
Covariance analysis. **(A, B, C)** *Top*: Structural covariance analysis centered on regions of cortical thickness group-differences. *Middle top:* Covariance patterns from each cluster were mapped in typically developing controls. *Middle bottom:* Group-wise differences in structural covariance was estimated at each vertex. Red/blue indicates increases/decreases in cortical thickness based on the relative contrast. Trend level differences are semi-transparent and significant regions for a cluster defining threshold of p<0.025, after cluster-level corrections for multiple corrections, are opaque. Black outlines indicate significant regions at corrected levels for cluster defining thresholds of p<0.05 *(A)* and p<0.025 *(B, C). Bottom*: Scatter plots illustrate group differences in covariance from seed regions to significant clusters.

**Table 3.** Structural covariance relationships described through correlation coefficients (β), standard error (SE) of coefficients, variance accounted for (R^2^), and average p-value of all covariance regions. Values correspond to the graphs in Figure 2 bottom and Figure S2.

| | $\beta$ | SE | $R^2$ | $p$ |
| --- | --- | --- | --- | --- |
| *IFG* |  |  |  |  |
| ANX | 0.80 | 0.08 | 0.79 | 0.008 |
| TD | 1.10 | 0.11 | 0.79 |  |
| SCARED | -0.01 | 0.007 | -1.75 | 0.086 |
| SRS | -0.002 | 0.002 | -0.83 | 0.041 |
| *CUN* |  |  |  |  |
| ASD | 0.73 | 0.12 | 0.58 | 1.06e-5 |
| TD | 1.04 | 0.15 | 0.64 |  |
| SCARED | -0.006 | 0.009 | -0.64 | 0.53 |
| SRS | -0.003 | 0.003 | -1.16 | 0.25 |

In TDC, the CUN showed strong covariance within bilateral occipital lobes as well as with the left precuneus and somatosensory cortex (**Figure 2C**). Compared to ANX, ASD was associated with weaker covariance of the CUN with bilateral ventrolateral prefrontal cortices (p_FWE_<0.001) (**Figure 2C, Table 3**). In TDC, the MTG exhibited strong covariance with somatosensory cortex (**Figure 2A**). The RSC showed a more locally confined pattern of covariance (**Figure S1**). The MTG and RSC clusters only showed marginal effects for an interaction when comparing ASD to TDC at the given cluster defining thresholds. However, at more exploratory cluster defining thresholds (p<0.05), we observed decreases in covariance in ASD compared to TDC between the MTG and association areas of the ipsilateral frontal (p_FWE_<0.001) and parietal (p_FWE_<0.001) cortices (**Figure 2A, Table 3**). Overall, when assessing covariance from regions of significant cortical thickness differences from between groups, we thus observed a shared decrease in covariance associated with the clinical groups compared to TDC as well as in ASD when compared to ANX.

#### 3.3 Phenotypic analyses

Analyses of behavioral phenotypes complemented the case-control results. At the behavioral level, analysis of variance indicated group differences in SRS-2 (F(2, 72)=35.11, p<0.001) and SCARED (F(2, 65)=8.84, p<0.001) (**Figure 3A)**. Post hoc Tukey tests showed graded increase in SRS-2 from TDC to ANX to ASD, while ASD and ANX did not differ on SCARED, but both were significantly greater than TDC. Second, considering regional cortical thickness measurements, higher SRS-2 scores were associated with increased cortical thickness in bilateral middle and superior temporal gyri (**Figure 3B**; t=2.50, p_FWE_<0.003), resembling the ASD-group difference analysis. Indeed, the t-statistic maps on the association of cortical with SRS-2 and ASD-vs-TDC (see *Figure 1*) exhibited strong correspondence (**Figure 3C**; r=0.71; p_spin_<0.001), suggesting convergent cortical substrates of phenotypic variables of autism risk and autism diagnosis (Bölte et al., 2011). Specificity for ASD was suggested, as the SRS-2 correlation map was weakly correlated with the ANX-vs-TDC contrast map (**Figure 3C**;r=0.30; p<0.001, spin permutation tests). Considering SCARED scores, we did not observe a significant association with cortical thickness after correction for multiple comparisons (**Figure 3B**); however, the uncorrected t-statistic map of the association between SCARED and cortical thickness was moderately correlated with the contrast map from the ANX-vs-TDC comparison (see *Figure 1;* **Figure 3C**;r=0.47; p_spin_<0.001) and weakly correlated with the ASD-vs-TDC contrast (**Figure 3C**;r=0.32, p_spin_<0.001).

**Figure 3.**
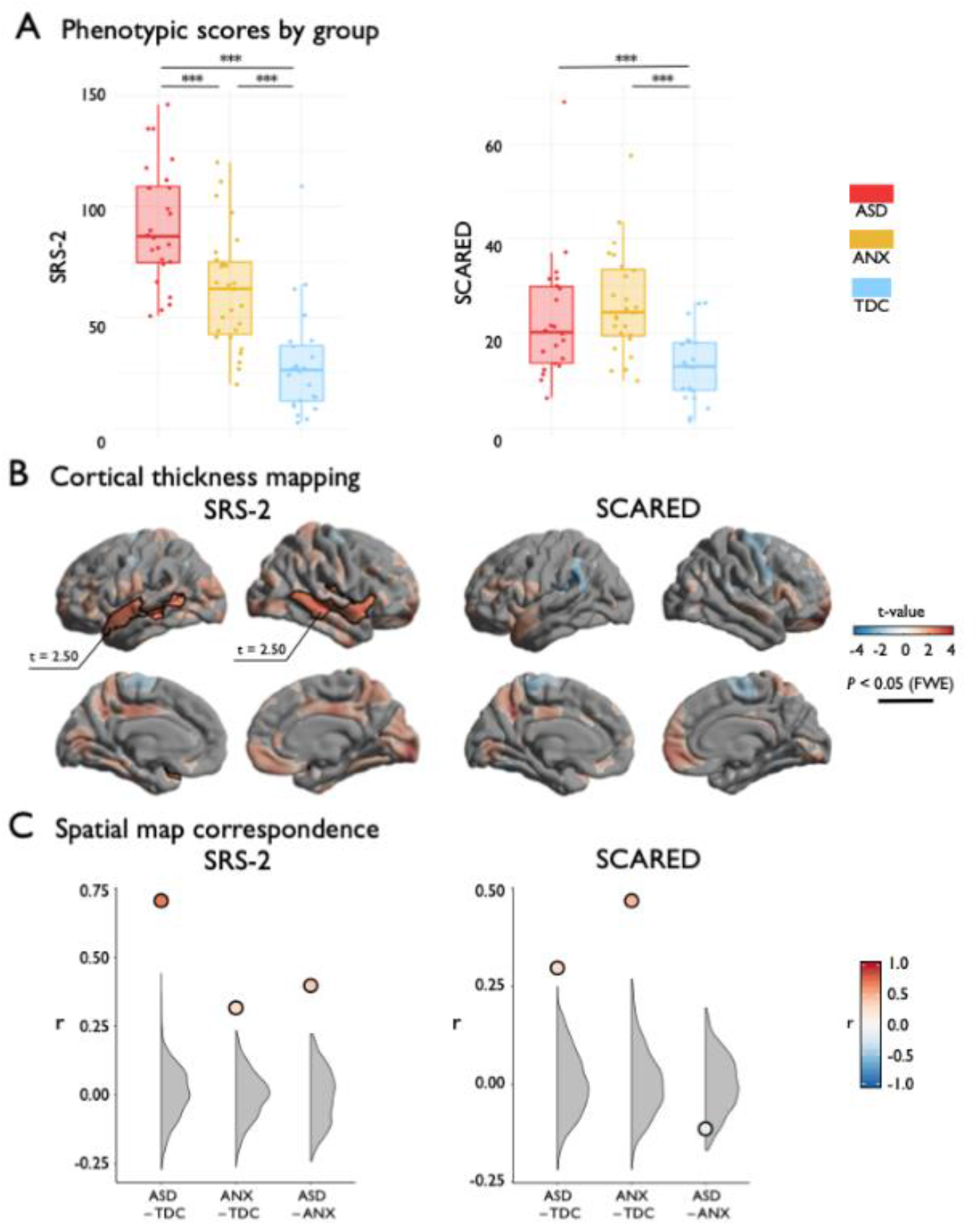
Phenotypic analysis. **(A)** SRS-2 and SCARED scores stratified by group. **(B)** Surface projection of the main effect of SRS-2 or SCARED on cortical thickness. Red/blue indicates increases/decreases in cortical thickness based on the relative contrast. Trend level differences are semi-transparent and significant regions after correction for multiple corrections outlined in black. **(C)** The spatial map correspondence between each group comparison and SRS-2 or SCARED compared to a null model distribution generated by the spin permutation method. *** p_spin_<0.001.

Finally, post-hoc analysis in regions of significant diagnosis-by-covariance modulations (**Figure S2**) indicated overall consistent findings, with high autism and anxiety symptoms being related to weaker covariance (more so for anxiety symptoms, p=0.086), suggesting symptom severity may in part recapitulates the network-level impact of the overall diagnostic category.

## 4. DISCUSSION

Our study aimed at identifying shared and distinct cortical substrates of primary diagnoses of ASD and ANX as well as phenotypic risk measures, using an integrated framework of regional and inter-regional structural MRI analyses. Our work benefitted from the first releases of the HBN (Alexander et al., 2017), an ongoing initiative aggregating and disseminating neuroimaging and phenotypic data in typically developing children as well as those with common neuropsychiatric conditions, such as ASD and ANX. Studying regional cortical morphology, we identified common and distinct morphological substrates associated with each condition. Although both groups presented with diffusely increased cortical thickness relative to controls, with highest overlaps in the inferior fronto-opercular regions, ASD was distinguished by additional thickness increases in temporal and posterior midline regions. Seed-based structural covariance analyses complemented these findings, showing common reductions in the morphological coupling between inferior prefrontal, lateral temporal, and temporo-parietal regions in both ANX and ASD compared to controls. Again, ASD was related to additional perturbances relative to ANX, specifically between midline parietal and prefrontal regions. Our case-control findings were complemented by phenotypic correlation analyses, lending a dimensional perspective on structural substrates of ASD and ANX risk. Behavioral phenotypes captured similar morphological patterns as group-level differences, particularly in the case of ASD. Collectively, our neuroanatomical findings support emerging transdiagnostic frameworks in highlighting common substrates of neurodevelopmental disorders, which are compounded by more extensive or severe abnormalities in more penetrative disorders, such as ASD.

Our regional analysis harnessed MRI-based cortical thickness measures, a reliable technique that has previously been applied to profile morphological variations across a broad spectrum of typical and atypical neurodevelopment (Raznahan et al., 2011). Surface-wide comparisons between ASD and TDC revealed increased cortical thickness in the former, with strongest effects in temporal and posterior midline regions, together with tendencies for prefrontal thickening, which is consistent with prior surface- and voxel-based data suggestive of diffuse grey matter increases in individuals with ASD (Bedford et al., 2019; Hong et al., 2017; Khundrakpam et al., 2017; Pereira et al., 2018; Raznahan et al., 2010; Scheel et al., 2011; Valk et al., 2015; Wallace et al., 2010). Although the ASD and TDC differences resembled those seen when comparing individuals with ANX to TDC, confirmed by spin tests that control for the shared spatial autocorrelations in two surface-based maps (Alexander-Bloch et al., 2018), ANX presented with a different distribution of anomalies with increased cortical thickness in fronto-insular-opercular regions while temporal and midline parietal were only visible at uncorrected significance level. While some lack in convergence can be attributed to limitations in sensitivity resulting from our modest sample size, our study benefitted from relatively strict inclusion criteria with respect to data availability and quality, together with the use of formal matching procedures that ensured similar age and sex distributions in our cohorts as well as a matched prevalence of ADHD comorbidity in ASD and ANX. This strategy may have increased specificity, possibly reflected in the divergence seen when directly comparing the clinical cohorts, revealing differential increases in cortical thickness in posterior midline regions in ASD compared to ANX.

Complementing regional morphological analysis, we harnessed an MRI covariance paradigm to examine large-scale coordination of structural networks. Covariance analyses have been extensively applied to study healthy brain development and to map perturbed structural networks across different neurological and neuropsychiatric diseases with early onset (Bernhardt et al., 2014a, b; Bethlehem et al., 2017; Park et al., 2018; Zuo et al., 2018). In line with ample data indicating reduced connectivity in ASD relative to TDC (Jung et al., 2019; McAlonan et al., 2005; Rudie et al., 2013; Sharda et al., 2014; Valk et al., 2015), we observed reduced covariance in ASD relative to TDC between temporal, prefrontal and parietal regions, suggestive of abnormal structural maturation of hetero- and transmodal association cortices. Although the link between covariance patterns and alternative *in vivo* metrics used in the connectomics field remains to be established (Gong et al., 2012), a prior study supports sensitivity to cortico-cortical maturation (Alexander-Bloch et al., 2013). As such, our findings in ASD likely indicate perturbations in large-scale maturational coordination, and may echo prior diffusion MRI findings showing atypical structural connectivity and microstructure in frontal, temporal and parietal white matter (Aoki et al., 2013; Rudie et al., 2013; Travers et al., 2012) as well as resting-state functional connectivity findings suggesting perturbed connectivity in higher-order systems such as the default mode network (Di Martino et al., 2013; Hong et al., 2019; Just et al., 2004; Jung et al., 2014; Kennedy and Courchescne et al., 2008; Picci et al., 2016), potentially by compromising the efficacy of long-range connections mediating higher-order and frequently self-generated functions (Di Martino et al., 2009). Similar to ASD, ANX also presented with reduced covariance of regions that were atypically thick relative to controls. Overall, the networks found to be decreased against controls in both cohorts were mainly anchored in lateral prefrontal, lateral temporal and temporo-parietal cortices, suggesting an that ASD and ANX may share a maturational perturbation of functional systems situated in higher-level association cortices.

In addition to the categorical case-control analyses, we examined the association of regional and inter-regional markers of cortical thickness with behavioral risk indices of ASD and ANX. Diagnostic and phenotypic associations closely overlapped in the case of ASD, but were less convergent in the case of ANX. This suggests that SRS-2 captures a relationship between cortical morphology and clinical phenotype that transcends diagnostic boundaries but is most severe in ASD. In contrast, SCARED scores were not significantly correlated with cortical thickness across the cohort. One previous study showed SCARED scores were related to decreased global cortical thickness in children, though excluded individuals with ASD from the analysis (Newman *et al.*, 2016). Together, these results suggest that the severity of anxiety symptoms, assessed by the SCARED questionnaire, has limited association with regional cortical morphology when taken across primary diagnoses. Both dimensional scales and ASD- and ANX-group comparisons displayed increased thickness in the inferior frontal gyrus, thus the area may mark severity of abnormal behaviour more generally. We also noted comparable differences in structural covariance of the inferior frontal gyrus between ANX and TDC groups and high-SCARED and low-SCARED. This observation once again suggests disruptions in structural networks that monitors salience in individuals with high anxiety traits, independent of a documented diagnosis (DiQuattro and Geng, 2011).

Our findings at the level of regional morphology and inter-regional covariance already lend support for the power of transdiagnostic approaches to unveil neurobiological factors that may play a role in the risk for prevalent mental health conditions such as ASD and ANX. The present study demonstrates that individuals with primary diagnoses of ASD and ANX exhibit distinctive patterns of cortical morphology, even though many individuals shared comorbid disorders. Shared comorbidities in ASD and ANX may explain the overlap of unthresholded comparisons with TDC. Such questions may be tested in the future as openly available datasets, such as the HBN repository, continue to grow and help to look for shared and unique effects beyond traditionally diagnostic categories.

## ACKNOWLEDGEMENTS

We would like to thank the Healthy Brain Network for providing the data for the present study. S.Y. received a scholarship from Natural Sciences and Engineering Research Council of Canada. R.A.I.B acknowledges research support by the Autism Research Trust and a British Academy Fellowship (PF2\180017). A.D. and M.M. were supported by NIMH (NIMH-105506 and NIMH-099059, respectively). C.P. received support from Fonds de la Recherche du Québec – Santé (FRQ-S). B.C.B. acknowledges support from CIHR (FDN-154298), SickKids Foundation (NI17-039), Natural Sciences and Engineering Research Council (NSERC; Discovery-1304413), Azrieli Center for Autism Research of the Montreal Neurological Institute (ACAR), and salary support from FRQS (Chercheur Boursier Junior 1). R.A.I.B and B.C.B were furthermore supported by an MNI-Cambridge collaboration grant.

